# Fndc3a (Fibronectin Domain Containing Protein 3A) influences median fin fold development and caudal fin regeneration in zebrafish by ECM alteration

**DOI:** 10.1101/386813

**Authors:** Daniel Liedtke, Melanie Orth, Michelle Meissler, Sinje Geuer, Sabine Knaup, Isabell Köblitz, Eva Klopocki

## Abstract

We investigated potential functions of Fndc3a during caudal fin development and regeneration in zebrafish. Reduced function interferes with correct epidermal cells structure and implies a role during vertebrate extremity development.

**Abstract:** Inherited genetic alterations are often found to be disease-causing factors of patient phenotypes. To unravel the molecular consequences of newly identified factors functional investigations *in vivo* are eminent. We investigated molecular functions of FNDC3A (Fibronectin Domain Containing Protein 3A; HUGO), a novel candidate gene for split-hand/foot malformations (SHFM) in humans, by utilizing zebrafish (*Danio rerio*) as a vertebrate model. Patients with congenital SHFM display prominent limb malformations, which are caused by disturbance of limb development due to defects in apical ectodermal ridge (AER) establishment and maintenance. Initial gene expression and protein localization studies clarified the presence of fndc3a in developing and regenerating fins of zebrafish. For functional studies we established a hypomorphic fndc3a mutant line (*fndc3a^wue1/wue1^*) via CRISPR/Cas9, exhibiting phenotypic malformations and changed gene expression patterns during early stages of median fin fold development. Furthermore, *fndc3a^wue1/wue1^* mutants display abnormal collagen localization, actinotrichia breakup and cellular defects in epidermal cells during caudal fin development. The observed effects are only temporary and later result in rather normal fin development in adults. In accordance with early fin development, proper caudal fin regeneration in adult *fndc3a^wue1/wue1^* mutants is hampered by interference with actinotrichia formation and epidermal cell abnormalities. Investigation of cellular matrix formation implied that loss of ECM structure is a common cause for both phenotypes. Our results thereby provide a molecular link between Fndc3a function during both developmental processes in zebrafish and foreshadow Fndc3a as a novel temporal regulator of epidermal cell properties during extremity development in vertebrates.

## Introduction

Patients suffering from SHFM display prominent limb malformations, lacking the central autopod rays, resulting in syndactyly, aplasia and/or hypoplasia of the phalanges, metacarpals and metatarsals, and median clefts of hands and feet (Duijf et al., 2003; Gurrieri and Everman, 2013). Several genetic loci and causes have been associated with this inherited disease, for example mutations in *TP63* (OMIM *603273), *DLX5* (OMIM *600028) or *DLX6* (OMIM *600030). A large number of animal experiments and cell culture studies further clarified the underlying genetic network, but still unresolved patient cases disclose the necessity to identify novel contributing molecular factors.

We identified *FNDC3A* as a new potential candidate gene in pathogenesis of human non-syndromic SHFM by exome sequencing of a consanguineous family from Syria with four affected individuals (unpublished data). FNDC3A consists of up to nine fibronectin type III domains, which are a common feature of a large number of extracellular proteins acting by modulation of different signaling pathways (Cheng et al., 2013; Zhu and Clark, 2014). *FNDC3A* has initially been described to be overexpressed in human odontoblasts (Carrouel et al., 2008). Functional experiments in *Symplastic spermatids (sys)* knockout mice indicated that FNDC3A is essential for cell adhesion between spermatids and Sertoli cells, resulting in sterile males (Obholz et al., 2006). Besides these two studies the developmental function of FNDC3A is still unknown and no association to extremity development or regeneration has been described yet. The purpose of this study was to investigate potential functions of Fndc3a during extremity development and regeneration in zebrafish.

Differences between tetrapod limb and fin development in ray-finned fishes (Actinopterygii) are obvious in anatomy and structure, but both are homologous appendages and thought to share common genetic features (Yano and Tamura, 2013). Also the development of pectoral fins in fish species is assumed to closely resemble extremity development in higher vertebrates, by sharing common molecular signals arising from a structure called the apical ectodermal ridge (AER) (Amaral and Schneider, 2017; Yano et al., 2012). Only recently differences between fin and limb AER have been reported and hint at a fin specific cellular process in fish species (Masselink et al., 2016; Yano et al., 2012). Even more eminent are developmental differences between paired fins (pectoral fins) and unpaired fins (caudal, anal and dorsal fins), as the establishment of the is achieved by evolutionary more distant processes (van den Boogaart et al., 2012). All unpaired fins arise from a common developmental precursor structure during larval stages called the median fin fold, which is exclusively found in larval teleosts. A wide number of molecular processes and distinct genes have been identified by several genetic screens in zebrafish revealing a complex network of factors necessary for correct median fin fold development (Carney et al., 2010; van Eeden et al., 1996). More than 30 years ago changes of epidermal cell shape and modulation of the extracellular matrix (ECM) have been described as one of the essential factors for correct median fin fold and caudal fin morphogenesis in zebrafish (Dane and Tucker, 1985). Later, signals from the WNT signaling pathway have been shown to be crucial for regulation of epithelial cell morphology by modulating laminin levels and thereby orchestrating correct patterning in growing fins (Nagendran et al., 2015). These early cellular steps of caudal fin development are prerequisites for subsequent processes; i.e. mesoderm cell migration, cell differentiation and normal fin growth. After initiation and pattern formation within the fin fold, further tissue differentiation results in the development of cartilage and bone, as skeletal elements in the later fin, and the gradual resorption of the median fin fold during juvenile stages (Parichy et al., 2009). Fin rays are formed by the assembly of actinotrichia and lepidotrichia in the ECM of epidermal cells (Grandel and Schulte-Merker, 1998; Zhang et al., 2010). While lepidotrichia are segmented and calcified bone rays, actinotrichia are non-calcified collagen fibers. Two collagens are essential for actinotrichia formation during fin development: Col2a1 and Col1a1 (Duran et al., 2011). In addition to the collagens actinodin proteins, encoded by the *and1* and *and2* genes, have been identified to be essential for actinotrichia formation (Zhang et al., 2010). During development actinotrichia fibers are normally formed at the fin tips, depicting characteristic brush-shaped structures.

Fin regeneration is a developmental process in adult fish sharing a number of conserved molecular mechanisms with extremity development at earlier stages of development (Iovine, 2007; Wehner and Weidinger, 2015). For both processes correct epidermal cell function, epithelial cell structure, and actinotrichia fibers are essential to correctly build all skeletal and mesenchymal fin elements (Mari-Beffa et al., 1989; Pfefferli and Jazwinska, 2015). Additional structural factors, especially ECM proteins like integrins and laminins, have been implied in regulating Wnt signaling during regeneration and in correct assembly of the teleost fin (Carney et al., 2010; Nagendran et al., 2015; Webb et al., 2007). Although a large number of involved “molecular players” have been described to date, not all factors necessary for correct ECM assembly in the regenerating caudal fin have been identified yet. We propose Fndc3a as a novel player in both fin development and regeneration.

## Results

The *FNDC3A* orthologous gene in zebrafish, *fndc3a*, is located on zebrafish chromosome 15 (ENSEMBL Zv9: 3,066, 162-3, 114,443 reverse strand; ENSDARG00000067569; ZFIN ID: ZDB-GENE-030131-7015) and encodes in 29 exons for a transcript of 3501 bp. The corresponding zebrafish 1166aa Fndc3a protein (ENSDART00000097261) consists of 4 transmembrane domains and 9 fibronectin type III domains. Two different predicted transcript variants have recently been reported for *fndc3a* (GenBank: XM_021466300.1, XM_021466301), encoding for proteins of 1247 and 1217aa. Both transcript variants are highly similar and differ only in a 30aa stretch at the N-terminus.

Phylogenetic and syntheny analyses showed that the *FNDC3A* gene is highly conserved throughout vertebrate evolution and orthologues are not duplicated in ray-finned fish species (data not shown). Amino acid alignments resulted in an up to 57% amino acid identity with 95% coverage, indicating a high level of conservation between human and zebrafish proteins. Two *fndc3a* paralogues can be identified in the zebrafish genome: *fndc3ba* (chromosome 2; ENSDARG00000078179; ZFIN ID: ZDB-GENE-070510-1) and *fndc3bb* (chromosome 24; ENSDARG00000062023; ZFIN ID: ZDB-GENE-070510-2). Both genes share highest sequence similarities with *FNDC3B* genes in other species and are dedicated in genomic analyses to an evolutionary distinct group when compared to *FNDC3* genes. All three gene family members have not been functionally investigated in zebrafish yet.

### Expression of *fndc3a* during early zebrafish development

To resolve the spatiotemporal expression of *fndc3a* during zebrafish development, we performed RNA in-situ hybridization experiments. *fndc3a* transcripts were detected spatially restricted to the tail bud region and the ventral fin fold mesenchyme from 14hpf onwards (hpf = hours post-fertilization; Fig. 1). Expression of *fndc3a* at later stages of embryonic development is mainly present in distinct brain regions; e.g. mid-hindbrain boundary, rhombencephalon, and mesencephalon; as well as in pectoral fins, in the notochord, in somites and in the caudal median fin fold (Fig. 1A and Fig. S8A). Restricted expression in the developing tail bud region at later stages (>16hpf) could be observed in the median fin fold, in Kupffer’s vesicle, and in cells of the chordo neural hinge region by longer staining times (Fig. 1B).

**Fig. 1:**
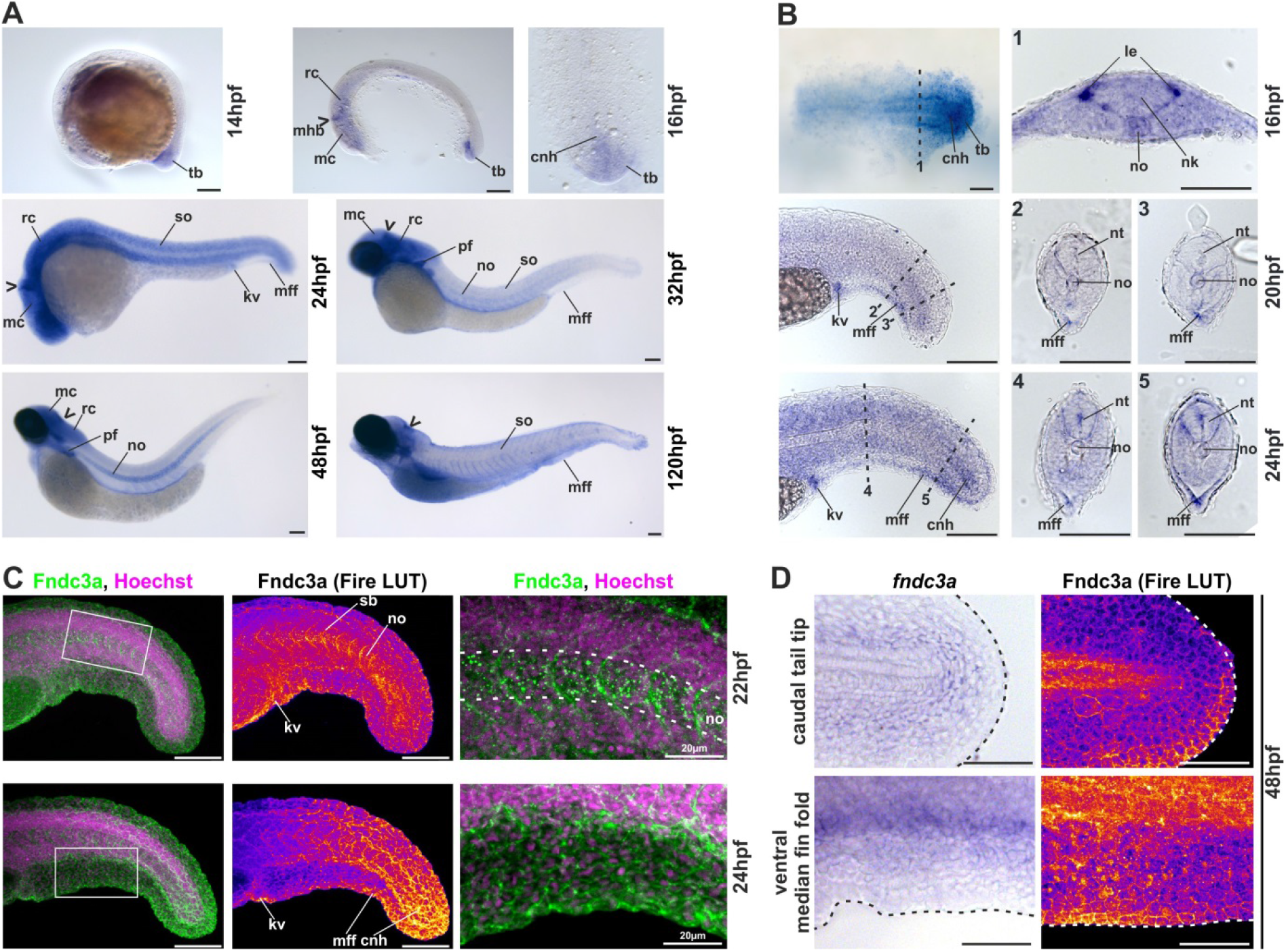
Localization of *fndc3a* RNA and protein during embryonic zebrafish development. (A and B) Expression of *fndc3a* is detected in the tail bud and the median fin fold from 14hpf onwards. During subsequent stages of zebrafish development *fndc3a* is expressed in pectoral fins, somites and notochord cells. Neuronal tissues express *fndc3a* starting from 16hpf in the midbrain-hindbrain boundary (marked by the chevron), the mesencephalon, and the rhombencephalon. (B) Investigation of long-time stained embryos and transversal sections indicated varying *fndc3a* expression in the tail bud region, at the lateral edge of the neural keel, at the ventral median fin fold, at Kupffer's vesicle, and in chordo neural hinge cells. (C) Detection of Fndc3a protein via immunofluorescence indicated similar regional localization as *fndc3a* mRNA in 22-24hpf embryos. Furthermore it showed intracellular accumulation of Fndc3a at notochord cells, at somite boundaries and epidermal cells at this stage. (D) Cellular localization of Fndc3a during median fin fold development at 48hpf corresponds to regions of mRNA expression and could be observed intracellular and at cell boundaries of epidermal cells. Dashed lines in B indicate planes of the corresponding numbered sections 1-5. Dashed lines in C indicate notochord boundary. Dashed lines in D indicate fin fold border. cnh: chordo neural hinge; kv: Kupffer's vesicle; le: lateral edge; mc: mesencephalon; mff: median fin fold; mhb: midbrain hindbrain boundary (marked with chevron); nk: neural keel; no: notochord; nt: neural tube; pf: pectoral fin; sb: somite boundary; so: somites; tb: tail bud; rc: rhombencephalon. Scale bars: 100μm, except higher magnification in C: 20pm.

Detection of Fndc3a protein localization was performed via immunofluorescence by application of a human FNDC3A antibody. This experiment showed similar regional localization of Fndc3a in Kupffer’s vesicle, the median fin fold region, and the caudal neuronal hinge during different stages of embryonic development consistent with RNA in-situ hybridization (24hpf: Fig. 1C and 48hpf: Fig.1D). The two most noticeable differences between RNA and protein localization were a spotted pattern of Fndc3a in notochord cells and in chevron shaped stripes between somite boundaries (higher magnification images in Fig. 1C). Moreover this experiment clarified the cellular localization of Fndc3a at the cell membrane of epidermal cells during early stages of median fin fold development (higher magnification images Fig. 1C and D).

### Generation of *fndc3a^wue1/wue1^* mutants

To investigate consequences of *fndc3a* loss in zebrafish, we established a *fndc3a^wue1/wue1^* mutant line via the CRISPR/Cas9 system (Hwang et al., 2013; Jao et al., 2013). sgRNAs were designed to target the evolutionary highly conserved third fibronectin III domain of Fndc3a (Fig. 2A), as incomplete genome information about the 5 end of *fndc3a* at the initiation of the project prevented the identification of a distinct start codon for sRNA targeting. For further analyses a zebrafish line with a 5bp substitution leading to a premature Stop codon in exon 13 (Zv9: ENSDARE00000690608) of *fndc3a* was chosen (*fndc3a^wue1^* line; ZFIN ID: ZDB-ALT-170417-3; Fig. S1A). The alteration was validated by sequencing of genomic DNA and cDNA (Fig. 2A; Fig. S1C) and could be detected continuously in subsequent inbred generations (data not shown). Off-site targets of the used *fndc3a* sgRNA were computationally predicted and investigated by sequencing the three most likely potential off-target sites. No off-site sequence alterations were detected in *fndc3a^wue1/+^ or fndc3a^wue1/+^* mutants (Fig. S1B).

**Fig. 2:**
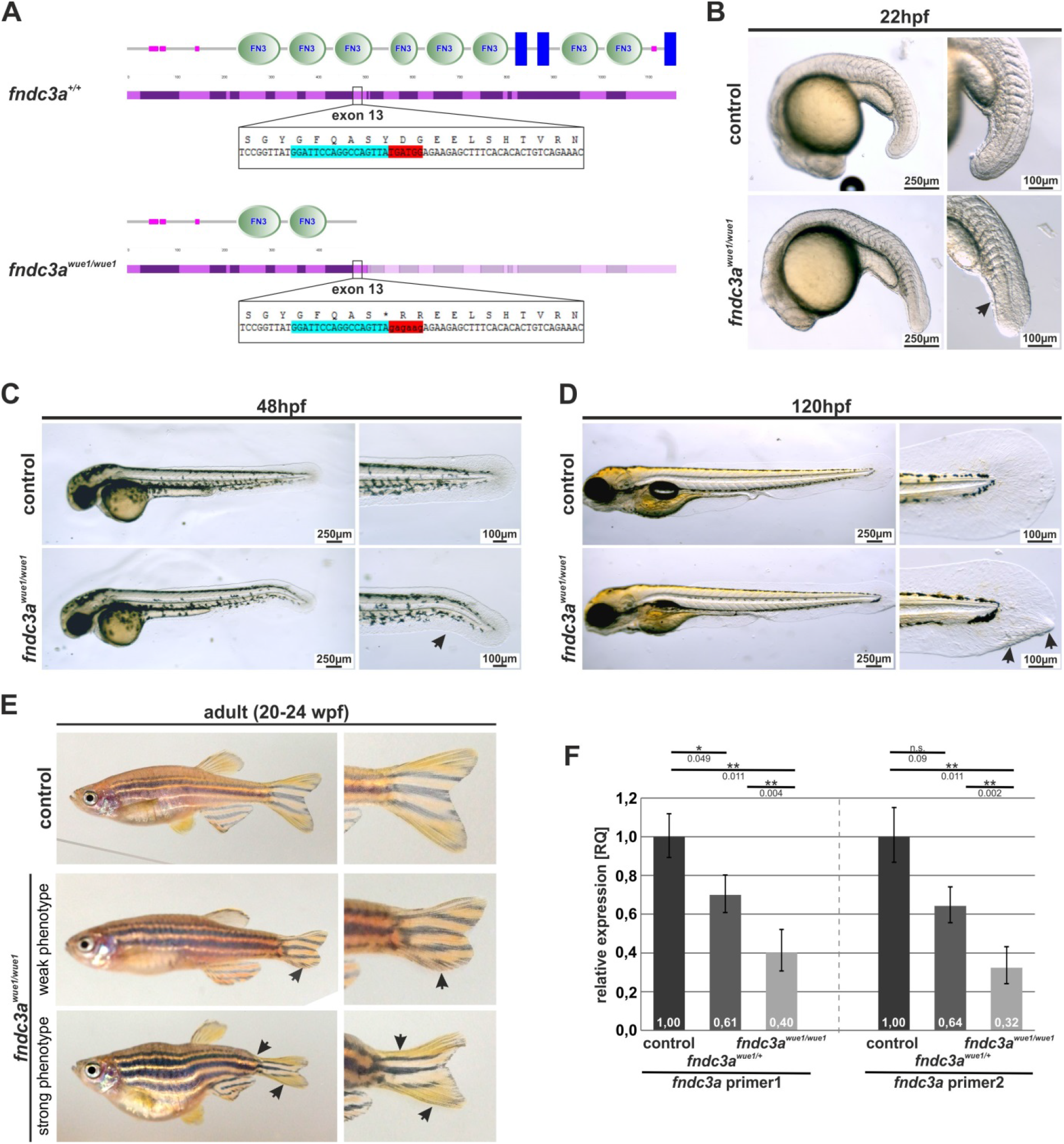
Generation and phenotype of *fndc3a^wue1,wue1^* zebrafish mutants. (A) The CRISPR/Cas9 system was used to target exon 13 in the zebrafish *fndc3a* gene coding for the third fibronectin type III domain (nucleotides marked in blue indicate sgRNA target sequence; nucleotides marked in red indicate the region of mutated sequence). (B-D) *fndc3a^wue1/wue1^* mutants show straightened tail buds (n=19/40), kinked tails (n= 27/100), and actinotrichia fiber aggregation during the first days of embryonic development (n= 9/41; indicated by arrows). (E) A fraction of adult *fndc3a^wue1/wue1^* mutants display weak (n= 15/71) to strong (n= 6/71) caudal fin phenotypes and tail malformations. (F) qPCR quantification of relative *fndc3a* expression levels in genotypic different groups of embryos indicated reduction of *fndc3a* transcripts in *fndc3a^wue1/+^* and *fndc3a^Nue1/Nue1^* (ΔΔCt calculation; comparison of biological triplicates for each genotype with twelve 24hpf embryos each; two independent *fndc3a* primer pairs; normalized against *AB* controls; *gapdh* and *ef1a1l1* expression were used as endogenous/housekeeping controls; significance levels and p-values of a 2-sided paired student t-test are given). Scale bars for embryo overview: 250pm; scale bars for tail magnifications: 100µm.

### Phenotypic investigation and validation of *fndc3a^wue1/wue1^* mutants

Initial visual investigation of *fndc3a^wue1/wue1^* mutants indicated a tail bud and median fin fold phenotype in homozygous embryos (Fig. 2B-D). A first indication of changed fin development could be observed 20-22hpf, as a number of *fndc3a^wue1/wue1^* embryos showed straightened tail buds and reduced ventral fin fold structures (47%, n=40; arrow in Fig. 2B). During subsequent stages of embryonic development *fndc3a^wue1/wue1^* mutant embryos developed kinked tails (27%, n= 100; 48hpf; Fig. 2C) and caudal fin fold malformations (22%, n= 41; 120hpf; arrows in Fig. 2D). The observed fin malformations in 120hpf embryos are sites of actinotrichia aggregation and show accumulation of apoptotic cells at this position (Fig. S2). Further assessment of temperature sensitivity of the *fndc3a^wue1/wue1^* phenotype revealed that raised temperatures result in an increased number and severity of the caudal fin phenotype (32°C incubation: 60%, n= 85; 28°C incubation: 48%, n= 40; 20°C incubation: 18%, n= 28; Fig. S3).

Besides the observed early median fin fold effects, *fndc3a^wue1/wue1^* juvenile and adult individuals in general showed normal development and behavior. The generated *fndc3a^wue1/wue1^* lines did not show reduction of fertility rates, which could have been suspected from published mouse knock-out lines (Fndc3a^Gt(RRP208)Byg^ or Fndc3a^sys^; Mouse Genome Database (MGD) at the Mouse Genome Informatics website, The Jackson Laboratory, Bar Harbor, Maine. http://www.informatics.jax.org; Obholz et al., 2006). Although, in approximately 30% of adult homozygous *fndc3a^wue1/wue1^* fish alterations of the caudal fin shape and the posterior body part could be visually detected (n=21/71). The observed phenotype ranged from minor fin shape changes in the majority of affected fish, up to axis shortening and stronger caudal fin deformations (Fig. 2E). We did not observe changes in phenotype severity or appearance rates in subsequent, homozygous generations, excluding a potential stronger maternal zygotic effect in *fndc3a^wue1/wue1^* or mitigation of the phenotype. The observation of variable adult phenotypes, incomplete phenotypic penetrance and temperature sensitivity of the embryonic phenotype led us to the assumption that *fndc3a^wue1/wue1^* mutants are hypomorphic. Further investigations of the *fndc3a^wue1/wue1^* mutant line via qPCR were performed to investigate potential nonsense mediated decay of *fndc3a* mRNAs. We quantified *fndc3a* mRNA expression in three independent embryo groups with two independent primer pairs (Fig. 2F; primer sequences are given in Table S1). Relative to *AB* controls *fndc3a* transcripts in *fndc3a^wue1/+^* and *fndc3a^wue1/wue1^* mutants were reduced to approximately 60% in heterozygotes and correspondingly to 30% relative expression in homozygotes. This experiment showed a reduction, but not a complete loss, of *fndc3a* mRNA in the *fndc3a^wue1/wue1^* mutants and thereby implies a potential residual function of Fndc3a in these mutants.

To achieve a full loss-of-function *fndc3a* phenotype two sgRNAs targeting exon 13 and 18 were designed and simultaneously used, resulting in a potential larger intragenic deletion or the introduction of several sequence alterations in the *fndc3a* locus. The corresponding transient phenotype indicated a more severe tail fin malformations in these fish (n=12/32) when compared to *fndc3a^wue1/wue1^* mutants (Fig. S4). Phenotypically the observed tail malformation defects showed strong caudal fin reduction and tail curling, similar to *smad5 (somitabun* or *piggytail*) (Mullins, 1996 #124;Hild, 1999 #123} or *bmp1a* zebrafish mutants (*frilly fins*) (Asharani et al., 2012). The double *fndc3a* CRISPR phenotype could not be maintained and next generation crossings did not result in a stable line, indicating, similar to *Symplastic spermatids (sys*) mice constricted fertility after complete loss or stronger reduction of Fndc3a function. However, this transient experiment supported our observations of the *fndc3a^wue1/wue1^* mutant and indicated a noticeable influence of Fndc3a function on caudal fin development.

### Reduction of Fndc3a function results in median fin fold defects

To elucidate the molecular causes underlying the early changes in caudal fin development of *fndc3a^wue1/wue1^* mutants we analyzed expression patterns of different, well established markers, expressed in the median fin fold tissues (Fig. 3). Expression of *fras1* is observed in the apical region of median fin fold (Carney et al., 2010; Gautier et al., 2008; Talbot et al., 2012). *hmcn1* expression is present in the epithelial cells of the apical fin fold, while expression of *hmcn2* can be detected in the fin fold epithelium and the fin mesenchyme (Carney et al., 2010; Feitosa et al., 2012). *bmp1a* expression is described in osteoblasts, fin mesenchyme cells, floor plate and hypochord cells (Asharani et al., 2012; Jasuja et al., 2006). *fbln1* is detected in presomitic mesoderm cells and in the dorsal neural rod (*Feitosa et al., 2012; Zhang et al., 1997*). In *fndc3a^wue1/wue1^* mutants expression of *fras1* and *hmcn1* were still present in the dorsal fin apical region 20 to 22hpf, but lost in the ventral region between Kupffer’s vesicle and the tail bud tip (number of affected *fndc3a^wue1/wue1^* embryos: *fras1*: 9/32; *hmcn1*: 11/26; Fig. 3A). Vice versa, expression patterns of *hmcn2, bmp1a*, and *fbln1* in this region were slightly changed and transverse sections indicated that ventral fin fold structures were lost (number of affected *fndc3a^wue1/wue1^* embryos: *hmcn2*: 24/43; *bmp1a*: 22/45; *fbln1*: 16/32). Expression of these three genes in the presomitic mesoderm was present in *fndc3a^wue1/wue1^* mutants but indicated convergence or even fusion of somites at ventral positions (indicated by arrows in Fig. 3A). The affected regions in the mutants correlated to the regions of *fndc3a* expression during the investigated stages (Fig. 1). Investigation of further mesodermal markers in *fndc3a^wue1/wue1^* mutants at this developmental stage additionally confirmed misplaced *myod* expression in ventral positions and indicated changes in chordo neural hinge cells by reduced *shha, ta(ntl)*, and *fgf8* expression (number of affected *fndc3a^wue1/wue1^* embryos: *myoD*: 12/22; *shha*: 7/17; *ta(ntl)*: 5/13; *fgf8a*: 5/13; Fig. S5), hinting at a structural or steric effect on tail development after Fndc3a reduction at this stage of development.

**Fig. 3:**
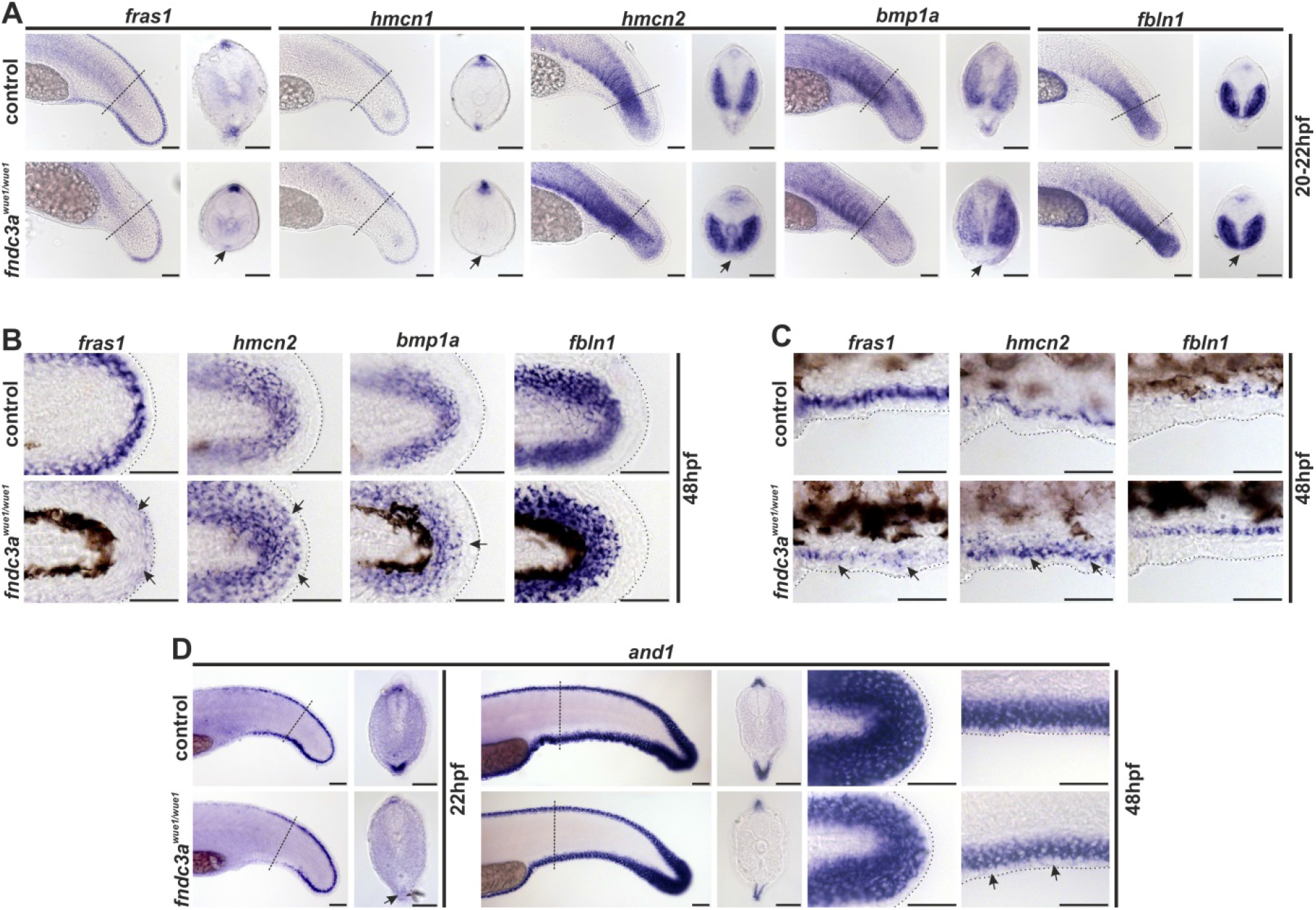
Normal development of the ventral median fin fold is altered in *fndc3a^wue1/wue1^* mutants. (A) Investigation of *fras1, hmcn1, hmcn2, bmp1a*, and *fbln1* expression in the tail bud 20-22hpf indicates loss of mesenchymal cells in the median fin fold in *fndc3a^Nue1ANue1^* mutants (arrows; *fras1*: 9/32; *hmcn1*: 11/26; *hmcn2*: 24/43; *bmp1a*: 22/45; *fbln1*: 16/32). (B) Expression of mesenchymal markers in posterior fins of 48hpf old embryos is slightly changed following loss of *fndc3a* (*fras1*: 10/24; *hmcn2*: 12/25; *bmp1a*: 8/21; *fbln1*: 9/24). (C) Ventral fin folds in 48hpf old embryos do not show loss of mesenchymal markers after *fndc3a^wue1/wue1^* mutation. (D) Expression of *and1* in *fndc3a^Nue1ANue1^* in the ventral fin fold is partly lost in 22hpf embryos (12/25), but recovers in 48hpf old embryos to a reduced expression level (control n=16; weaker expression in *fndc3a^wue1/wue1^* embryos n=10/18). Dashed lines in A and D indicate planes of shown sections. Dashed lines in B and C indicate fin boundaries. Scale bars: 50µm

Analyses in 48hpf mutant embryos showed reduced expression of *fras1* in *fndc3a^wue1/wue1^* mutants, but a slight increase of *hmnc2, bmp1a* and *fbln1* expressing cells in the posterior fin tip (number of affected *fndc3a^wue1/wue1^* embryos: *fras1*: 10/24; *hmcn2*: 12/25; *bmp1a*: 8/21; *fbln1*: 9/24; Fig. 3B). In the same set of embryos expressional changes of *fras1, hmcn2*, and *fbln1* in the ventral fin fold, posterior to the proctodeum were less evident, and indicated only a slight reduction of *fras1* expression and a slightly broader *hmcn2* expression (Fig. 3C). Similar to the investigated markers the ventral expression domain of *and1*, an essential factor for actinotrichia formation (Duran et al., 2011), was also different in 22hpf *fndc3a^wue1/wue1^* embryos (number of affected *fndc3a^wue1/wue1^* embryos: 12/25; Fig. 3D). *and1* expression was recovered in fin folds of 48hpf embryos, but seemed to be slightly weaker in comparison to controls (control n=16; weaker expression in *fndc3a^wue1/wue1^* embryos n=10/18). In general, the gene expression analyses indicate that *fndc3a^wue1/wue1^* embryos started to recover normal gene expression patterns in median fin fold cells and developed only temporal defects during early median fin fold development. Most prominently apical cells of the median fin fold were influenced by reduced levels of Fndc3a during the first 2 days of zebrafish development.

### Morpholino knockdown and rescue experiments support specific role of Fndc3a in median fin fold development

Validation of *fndc3a^wue1/wue1^* knockdown specific effects on median fin fold development in mutants was done by Morpholino knockdown and rescue experiments (Fig. S6 and S7). Phenotypic investigation of *fndc3a* morphants further clarified, that similar to *fndc3a^wue1/wue1^* mutant embryos, median fin fold malformations were induced in a dosage dependent manner in the first 48h of development (AB control embryos with tail phenotype 24hpf injected with 0.1mM *fndc3a* MO: 0/35; 0.25mM *fndc3a* MO: 17/96; 0.5mM *fndc3a* MO: 29/56; control: 2/348; Fig. S6B and C). Notably, injection of *fndc3a* Morpholino into *fndc3a^wue1/wue1^* embryos resulted in an enhanced number of embryos showing tail phenotypes and supports the hypothesis of a hypomorphic mutation (*fndc3a^wue1/wue1^* embryos with tail phenotype 24hpf injected with 0.1mM *fndc3a* MO: 12/36; 0.25mM *fndc3a* MO: 42/81; 0.5mM *fndc3a* MO: 17/18 with high mortality rates; control: 51/207; Fig. S6B and C). Loss of median fin fold markers *fras1* and *hmcn1* and changes in *hmcn2* expression in ventral fin folds were observed in *fndc3a* morphants and further confirm the observations in *fndc3a^wue1/wue1^* mutants (summarized altered marker gene expression in control: 0/57; *fndc3a^wue1/wue1^*: 30/58; 0.25nM *fndc3a* MO injection in AB: 14/28; *fndc3a* 0.25nM MO injection in *fndc3a^wue1/wue1^*: 12/12; Fig. S6D).

Rescue and overexpression experiments were performed by injection of full-length human *FNDC3A* RNA (Fig. S7). These experiments indicated a rescue of the tail phenotype in *fndc3a^wue1/wue1^* mutants after moderate RNA supplementation (tail phenotype in uninjected *fndc3a^wue1/wue1^*: 51/207; *fndc3a^wue1/wue1^* rescue with 25ng/μl *FNDC3A* RNA: 9/107; *fndc3a^wue1/wue1^* rescue with 50ng/µl *FNDC3A* RNA: 64/94; Fig. S7A and B). In addition, regain of *fras1* and *hmcn1* and *hmcn2* marker gene expression was detected in ventral median fin folds of rescued embryos (summarized altered marker gene expression in control: 0/20; *fndc3a^wue1/wue1^*: 25/55; *fndc3a^wue1/wue1^* rescue with 25 mg/μl *FNDC3A* RNA: 4/80; Fig. S7C). Injection of *FNDC3A* RNA into *AB* control embryos did rarely result in tail malformations (tail phenotype in control: 2/345; 25ng/μl *FNDC3A* RNA: 4/36; 50ng/μl *FNDC3A* RNA: 7/93; Fig. S7B). These experiments indicate a narrow threshold level for Fndc3a to fulfill its function in early median fin fold cells and support the observed *fndc3a^wue1/wue1^* phenotype.

### *fndc3a^wue1/wue1^* mutants show actinotrichia breakdown and basal epidermal cell defects

Our observations in homozygous mutants and in morphants implied a transient function of Fndc3a during early median fin fold development resulting in caudal fin malformation at later stages of development. Localization of *fndc3a* mRNA expression and Fndc3a protein during these stages further suggests that the observed effects are directly or indirectly linked to epidermal cells or associated structures. To clarify this question and to investigate structural defects causing the observed developmental phenotype we at first investigated actinotrichia formation in *fndc3a^wue1/wue1^* mutants.

Actinotrichia are a fish-specific structural element consisting of collagen fibers (Col1a1 and Col2a1) and actinodin proteins (Duran et al., 2011; Zhang et al., 2010). Characteristically transverse striation of actinotrichia can be observed in several fish species via electron microscopy and suggests that these fibers are giant collagen fibrils of hyperpolymerized collagen (Montes et al., 1982). Actinotrichia were visualized in *fndc3a^wue1/wue1^* mutants either by differential interference contrast (DIC) microscopy (Fig. 4A) or by immunofluorescent staining of Col2a, a structural component of actinotrichia ((Duran et al., 2011); Fig. 4B). Actinotrichia fibers in *fndc3a^wue1/wue1^* mutants were still present, but displayed obvious structural alterations and signs of breakdown in the ventral caudal fin (control: 0/10; *fndc3a^wuo1/wuc1^*: 7/12). While control fish at 52hpf showed radiant symmetrical arrangement of Col2a in the actinotrichia fibers of the developing caudal fin, *fndc3a^wuo1/wuc1^* embryos partly lack these structures and depicted unstructured, crumbled collagen fibers in the fin mesenchyme (control: 0/7; *fndc3a^wue1/wue1^*:12/14). High levels of remaining Col2a in *fndc3a^wue1/wue1^* mutants could be detected in apical cells at the fin border (arrows in Fig.4B), which were also visible as distinct cells in the DIC microscopy (Fig. 4A).

**Fig. 4:**
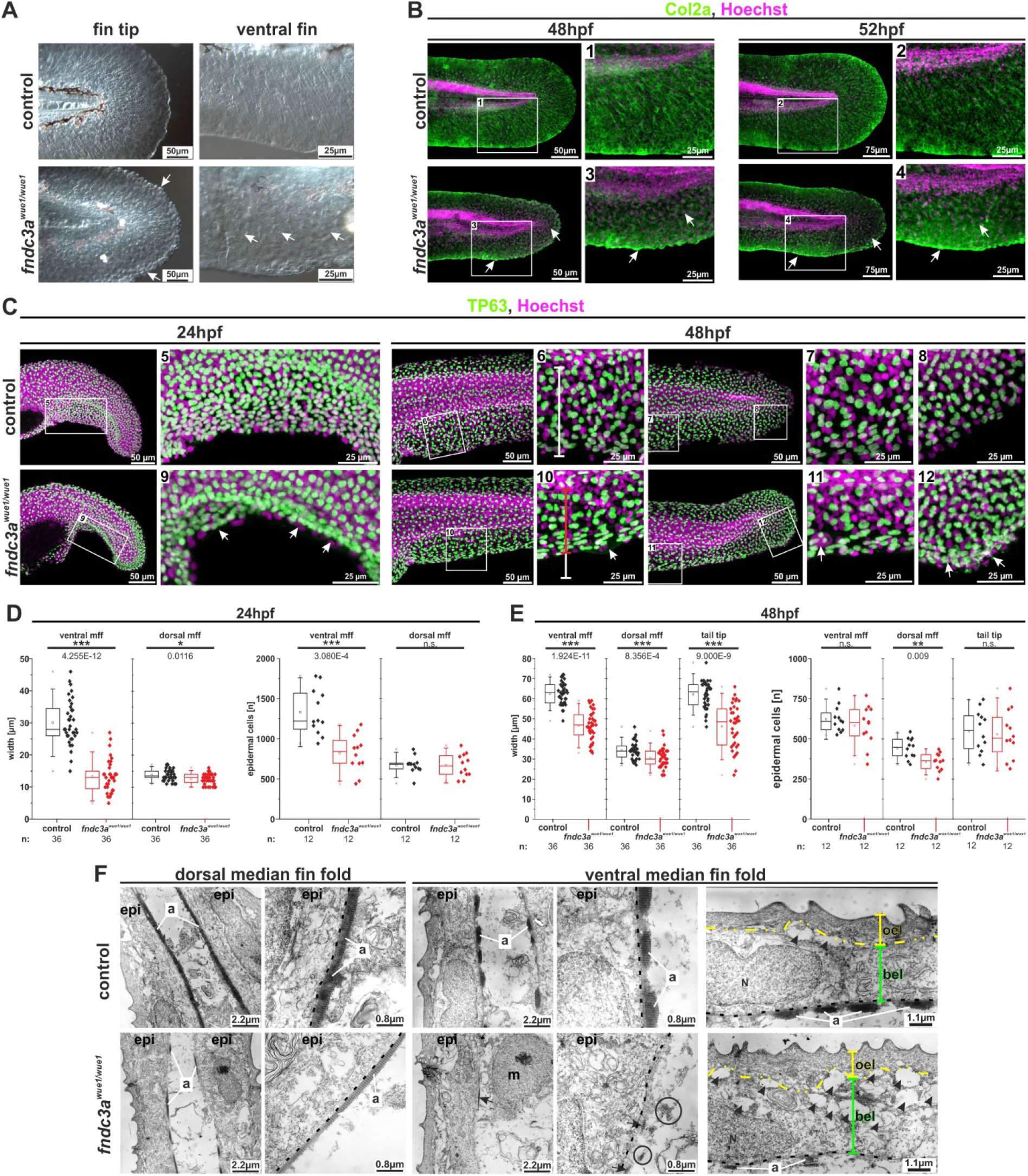
The *fndc3a^wue1/wue1^* mutation results in structural defects in epidermal cells during fin development. Visualization of actinotrichia either by differential interference contrast microscopy (A) or by immunofluorescence staining of Col2a (B) showed loss of mature actinotrichia in *fndc3a^wue1/wue1^* mutants 48 to 52 hpf (number of affected embryos: control = 0/17; *fndc3a^Nue1ANue1^* = 19/26; white arrows indicate lost actinotrichia and Col2a accumulation in apical cells of the median fin fold). (C) Investigation of TP63 positive cells in in *fndc3a^wue1/wue1^* mutants showed reduced width and reduced epidermal cell number in the ventral median fin fold in *fndc3a^wue1/wue1^* mutants. (D, E) Quantification of median fin fold width and TP63 positive cells in 24 and 48hpf embryos (Mann-Whitney U test; p<0.05; two-tailed, U values are indicated for significant changes). (F) Ultrastructural analyses of dorsal and ventral fins of control (upper row) or *fndc3a^wue1/wue1^* mutants (lower row) reveal breakdown of actinotrichia fibers and cellular malformations in cells of the basal epidermal layer in the ventral fin folds of 52hpf old embryos (arrows indicate site of remaining actinotrichia, circles indicate misplaced fibers; dashed grey lines indicate the basal membrane; yellow lines indicate outer epidermal cells and green lines indicate basal epidermal cells; Black arrowheads indicate cavities). a: actinotrichia; bel: basal epidermal layer; epi: epidermis; mff: median fin fold; m: mesenchymal cell; N: nucleus; oel: outer epidermal layer.

Potential cellular reasons for actinotrichia breakdown and Col2 misallocation in the median fin fold can be numerous. Based on the expression timing of *fndc3a* during median fin fold development and Fndc3a localization we assumed that epidermal cells might be influenced by reduced function of Fndc3a and that the normal cellular organization in the developing fin is lost (Dane and Tucker, 1985; Duran et al., 2011). To assess the potential effects of *fndc3a* mutation on epidermal cells during median fin fold development we performed staining for TP63 in *fndc3a^wue1/wue1^* mutants (Lee and Kimelman, 2002) (Fig. 4C). In 22-24hpf embryos epidermal cells of *fndc3a^wue1/wue1^* mutants showed lack of ventral fin fold structures and reduced median fin fold width at this position (arrows in Fig. 4C9; quantification in Fig. 4D). Along with this observation, a reduced number of epidermal cells could be detected at this stage in the ventral fin fold. At 48hpf *fndc3a^wue1/wue1^* mutants developed outgrowing ventral median folds. Although these were still shorter in length they did not display a significant lower number of epidermal cells in comparison to control embryos (Fig. 4D10 and D11; quantification in Fig. 4E), indicating cellular recovery of the median fin fold structure. Additionally, *fndc3a^wue1/wue1^* mutants displayed aggregation of epidermal cells at the fin fold border, which were not observed in control embryos (Fig. 4D12).

To further clarify the cellular consequences of Fndc3a reduction in epidermal cells of the median fin fold, we performed Transmission Electron Microscopy (TEM). Control or *fndc3a^wue1/+^* embryos showed normal arrangement of anatomical fin structures: the outer epithelial layer (oel), the basal epidermal layer (bel), the basal membrane (bm), and actinotrichia (a) in dorsal and ventral trunk fin folds (Fig. 4F). Actinotrichia fibers are normally attached to the basal membrane and display prominent stratification (van den Boogaart et al., 2012). *fndc3a^wue1/wue1^* mutant embryos showed an almost complete loss of actinotrichia in ventral fin folds, while in dorsal fin folds of the same individuals actinotrichia were only slightly reduced but still present (lower row Fig. 4F). Small remains of actinotrichia, showing characteristic stratification, were present in ventral fin folds (arrows Fig. 4F). But also misplaced collagen fibers not attached to the basal membrane could be detected (circles Fig. 4F). The ventral median fin fold of *fndc3a^wue1/wue1^* mutants displayed the presence of mesenchymal cells, the outer and basal epidermal cell layer (Fig. 4F). But the sub-cellular structure of the basal epidermal layer was altered. It exhibited cavities between the outer and basal epidermal layers (marked by arrow heads) and displayed changes in the ECM surrounding these cells. This observation strongly implies that the disruption of actinotrichia fibers and the observed effects on TP63 positive cells after Fndc3a reduction are due to cellular defects in the basal epidermal cell layer.

Contemporaneous to the caudal median fin fold, pectoral fins are formed during embryonic development and share common structural components and genetic factors with caudal fins (Iovine, 2007). RNA in-situ hybridization and immunofluorescence indicated *fndc3a* expression and localization during the first days of embryonic development in epidermal cells of pectoral fins (Fig. 1; Fig. S8A). A potential effect of *fndc3a* reduction on pectoral fin development was investigated by phenotypic observation (Fig. S8B) and Col2a as well as TP63 immunofluorescence staining (Fig. S8C). These experiments indicated no alterations in pectoral fin development in *fndc3a^wue1/wue1^* during embryogenesis. Likewise adult *fndc3a^wue1/wue1^* mutants and transient double fndc3a CRISPR injected fish did not show changed pectoral fin morphology (data not shown) and indicated no or only a minor role of Fndc3a in pectoral fin development.

### Fin regeneration is only temporally influenced in *fndc3a^wue1/wue1^* mutants

Besides their function during fin development, actinotrichia and epidermal cells possess essential roles during the regeneration of adult caudal fins after amputation (Duran et al., 2011; Santamaria and Becerra, 1991). Similar to their role in fin development Col2a and Col1a were identified to be necessary factors of actinotrichia formation during this process (Duran et al., 2015), implying comparable cellular processes during fin development and regeneration. We therefore hypothesized that reduction of Fndc3a function might also interfere with actinotrichia formation and basal epidermal cells during fin regeneration.

To test this hypothesis, we performed regeneration experiments on adult caudal fins with control and *fndc3a^wue1/wue1^* fish (Fig. 5 and Fig. S9). Two remarkable observations were made: First, regenerates looked opaque, disorganized and tubercular extensions attached to the epidermal layer of the fin regenerates were eminent between 4dpa and 6dpa (days post amputation) in *fndc3a^wue1/wue1^* mutants (arrowheads in Fig. 5A; Fig. S9B and D; abnormal regenerate phenotypes 6dpa: control: 3/19; *fndc3a*^wue1/wue1^: 13/19). Subsequent histological investigation via H&E staining on sections of these regenerates confirmed detached or loosely attached cells in the outer epidermal layer of the *fndc3a^wue1/wue1^* regenerates. These cells were additionally investigated by immunofluorescence and depicted high levels of Col2a at the regenerative front (arrows in Fig. 5B). TEM analysis further clarified that these cells were still attached to the epidermal cell layer and incorporate electronic dense material in their Golgi apparatus and in intracellular vesicles (Fig. 5E). In accordance with this observation, *fndc3a* expression in fin regenerates could be detected in the distal wound blastema at 4dpa and 6dpa (Fig. S9C). Localization of Fndc3a protein in regenerates was confirmed by immunofluorescence in the epidermal cell layers of regenerates (Fig. 5C). Second, actinotrichia fibers at the tip of 4dpa regenerates looked disorganized and clumped in *fndc3a^wue1/wue1^* mutants (arrows in Fig. 5A). In 4dpa regenerates of control individuals TEM analysis of formed actinotrichia fibers showed bundles of stratified actin fibers in close proximity to the basal membrane cells (Fig.5F). In contrast, *fndc3a^wue1/wue1^* mutants lack these prominent compact actinotrichia fibers and only depicted loose filaments adjacent to the membrane layer. Subsequent investigation of epidermal cells by TP63 staining in regenerates 4dpa showed interference with normal regenerate structure and disturbance of epidermal cells in *fndc3a^wue1/wue1^* mutant regenerates (Fig. 5D). This indicates altered epidermal organization as a potential reason for the observed effects in regenerates after reduced Fndc3a function, similar to the processes observed in the median fin fold during early development.

**Fig. 5:**
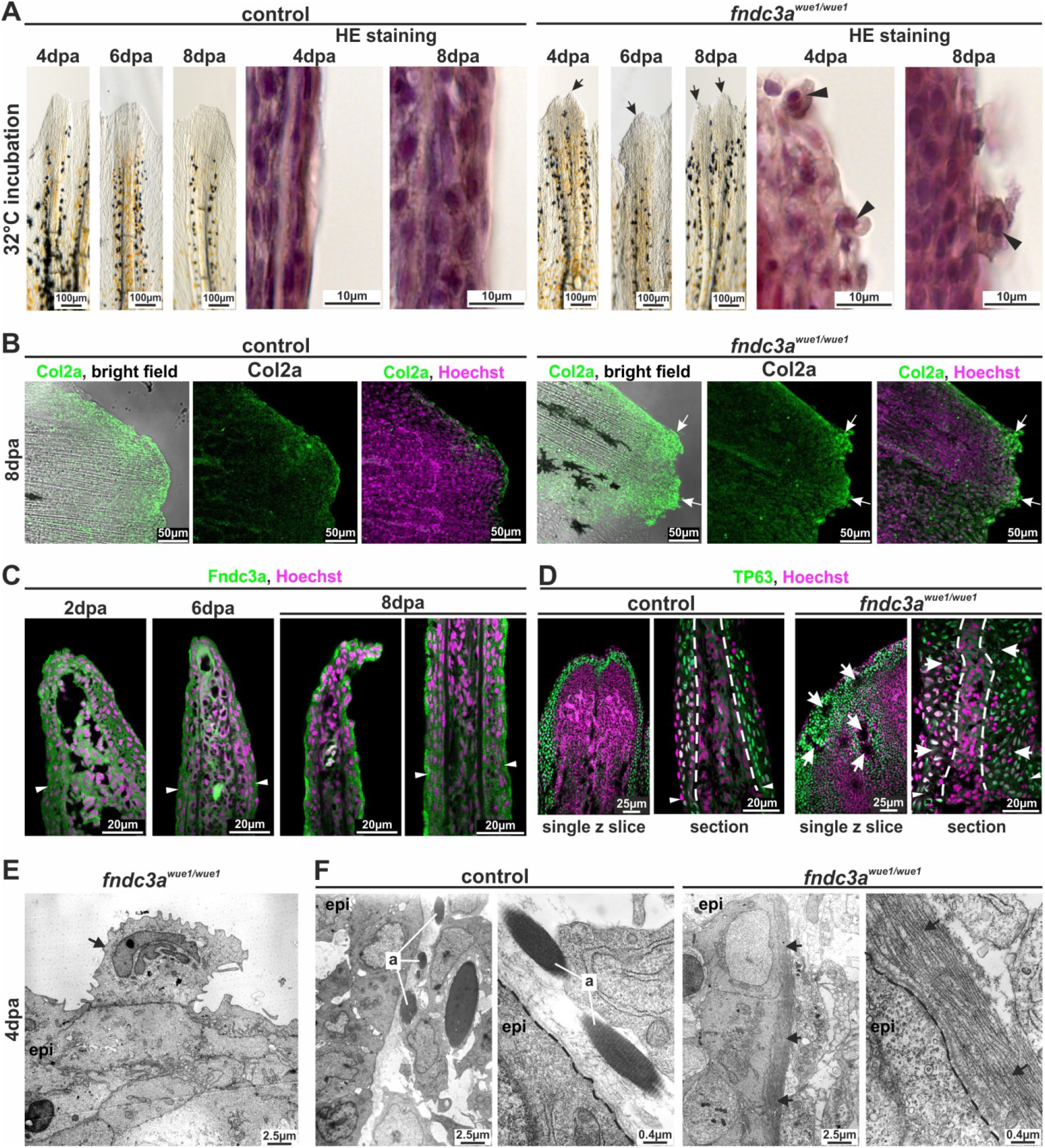
Interference with Fndc3a function during fin regeneration results in epidermal cells defects. (A) Phenotypical and histological investigations of fin regeneration in *fndc3a^wue1/wue1^* mutants indicate abnormal regeneration by showing unorganized fin borders at the regenerative front (arrows) and aberrant epidermal cells (arrowheads; abnormal regenerate phenotypes 6dpa: control: 3/19; *fndc3a*^Nue1ANue1^: 13/19). (B) Col2a protein localization visualized via immunofluorescence on 8dpa regenerates shows accumulation of collagen in these abnormal cells of *fndc3a^wue1/wue1^* mutants (white arrows; n=6 for each group). (C) Localization of Fndc3a in regenerates can be detected in epidermal cell layers (white arrowheads indicate amputation site). (D) TP63 staining in regenerates of *fndc3a^wue1/wue1^* mutants 4dpa indicates disorganization of epidermal cells (white arrows; white arrowheads indicate amputation site; n=4 for each group). (E) TEM analysis showed accumulation of electron dense material in the Golgi apparatus of abnormal cells attached to the epidermal layer (arrow). (F) TEM analyses of actinotrichia in fin regenerates 4dpa reveal loss of these fibers located near the basal membrane (arrows) in *fndc3a* depleted fish. a: actinotrichia; epi: epidermis; dashed lines indicate basal membrane.

Similar to the temperature dependency of the hypomorphic *fndc3a^wue1/wue1^* phenotype during fin development, the observed effects during fin regeneration could be enhanced by keeping fish at a raised temperature of 32°C during the phase of regeneration (incubation at 24°C in Fig. S9A and B; incubation at 32°C in Fig 5A and Fig. S9D). Although prominent cellular abnormalities were detected during the first days past amputation (dpa), the investigated control and mutant fish showed no significant differences in overall tail length growth in the first 10dpa and only at 6dpa a difference in regenerate length was detected (Fig. S10). All fins grew normally to their former size after ~14dpa (data not shown), but small changes in fin morphology, e.g. cooped fin rays and loss of intersegmental tissues, could be observed in *fndc3a^wue1/wue1^* mutants 6 weeks past amputation (wpa; control: 1/10; *fndc3a^wue1/wue1^*: 8/10; Fig. S9E). Our regeneration experiments hint to rather minor and temporal effects of Fndc3a on regeneration during initiation and the first few days of regeneration and point to a potential compensatory mechanism or residual function of Fndc3a in the *fndc3a^wue1/wue1^* mutants.

### *fndc3a^wue1/wue1^* mutants display disorganized cellular arrangement in median fin folds and in caudal fin regenerates

Correct assembly of actinotrichia and correct caudal fin morphology are greatly dependent on the extracellular composition and the cell shape of surrounding epidermal cells during development and regeneration. These cellular characteristics within the median fin fold determine correct signaling, e.g. by Wnt signals, and cell behavior (Wehner et al., 2014; Wehner and Weidinger, 2015). Fibronectin domain containing proteins like Fndc3a have been linked to functions during ECM assembly and maintenance (Henderson et al., 2011). Thus, we assume that Fndc3a function during median fin fold development and caudal fin regeneration is provoked by cell shape or by ECM alterations in epidermal cells.

To follow-up on this assumption we analyzed cell membrane structure and the ECM in the median fin fold of control and *fndc3a^wue1/wue1^* mutants by investigating F-actin (Phalloidin staining) or β-catenin localization (Fig 6). Cell boundaries of control embryos displayed a dense, stereotypical assembly of epidermal cells in the ventral median fin fold at 22hpf (Fig. 6A). Reduction of Fndc3a resulted in clustering of cells, altered cell shapes of ventral median fin fold epidermal cells (white arrows Fig. 6A) and appearance of cavities within the fin folds (white arrowheads and white dashed lines Fig. 6A). The cavities are zones within the tissue showing no F-actin and no nuclear staining, suggesting cell free spaces within the median fin fold. Further cellular observations were made by investigation of β-catenin localization at similar developmental stages (Fig. 6B). Homogeneous membranous localization was detected in cells of the ventral median fin fold of control embryos 24hpf and nuclear localization of β-catenin was detected in epithelial cells at the fin border (arrowheads in Fig. 6B)(Nagendran et al., 2015). In *fndc3a^wue1/wue1^* mutants uniform localization of β-catenin at the cell membrane of epithelial cells was impaired and instead appeared as accumulations of the protein in a speckled fashion within the cytoplasm (white arrows in Fig. 6B). Nuclear localization of β-catenin was not completely abolished in *fndc3a^wue1/wue1^* mutants and was clearly detected in epithelial cells at the apical fin border, indicating maintenance of a Wnt gradient in these mutants (grey arrowheads in Fig. 6B). These experiments suggest a partial loss of epidermal cellular structure and loss of adhesion within the median fin fold of *fndc3a^wue1/wue1^* mutants during embryonic development.

Besides the effects on median fin fold development we additionally investigated potential ECM changes in regenerates of *fndc3a^wue1/wue1^* mutants by F-actin (Fig. 6C and 6D) and β-catenin staining (Fig. 6E and 6F). Comparison between controls and *fndc3a^wue1/wue1^* mutants clarified that similar distinctive features, i.e. altered cell matrix and appearance of cavities within the tissue, were also observed in regenerates between 2 and 8dpa. F-actin depicted altered regenerate border shapes of *fndc3a^wuo1/wue1^* mutants (dashed white line Fig. 6C) and appearance of regions showing cellular alterations (white arrows Fig. 6C). These obviously fragmented structures within the blastema appeared 2dpa and could be detected until 6dpa. High resolution microscopy clarified, that fragmented structures in 2dpa regenerates are cavities within the blastema (white arrowheads Fig. 6D). Cellular alterations were also detected in cells at the regenerative front of *fndc3a^wue1/wue1^* mutants at 4 to 8dpa by β-catenin immunostaining (Fig. 6E). Irregular blastema borders 4dpa were detected distinctly with the same staining (white arrows and close-up pictures in Fig. 6E). High resolution imaging also revealed β-catenin negative, detached cells outside of the regenerate (white arrowheads in Fig. 6F) and cavities (white arrows in Fig. 6F). Later stages of caudal regeneration did not seem to be compromised by reduced Fndc3a level, as fin regenerates of *fndc3a^wue1/wue1^* mutants were able to grow to similar fin lengths as control fish at 10dpa (Fig. S10). Our observations thereby indicate a partial loss of cellular structure and loss of adhesion within the blastema during early stages of caudal fin regeneration in *fndc3a^wue1/wue1^* mutants.

**Fig. 6:**
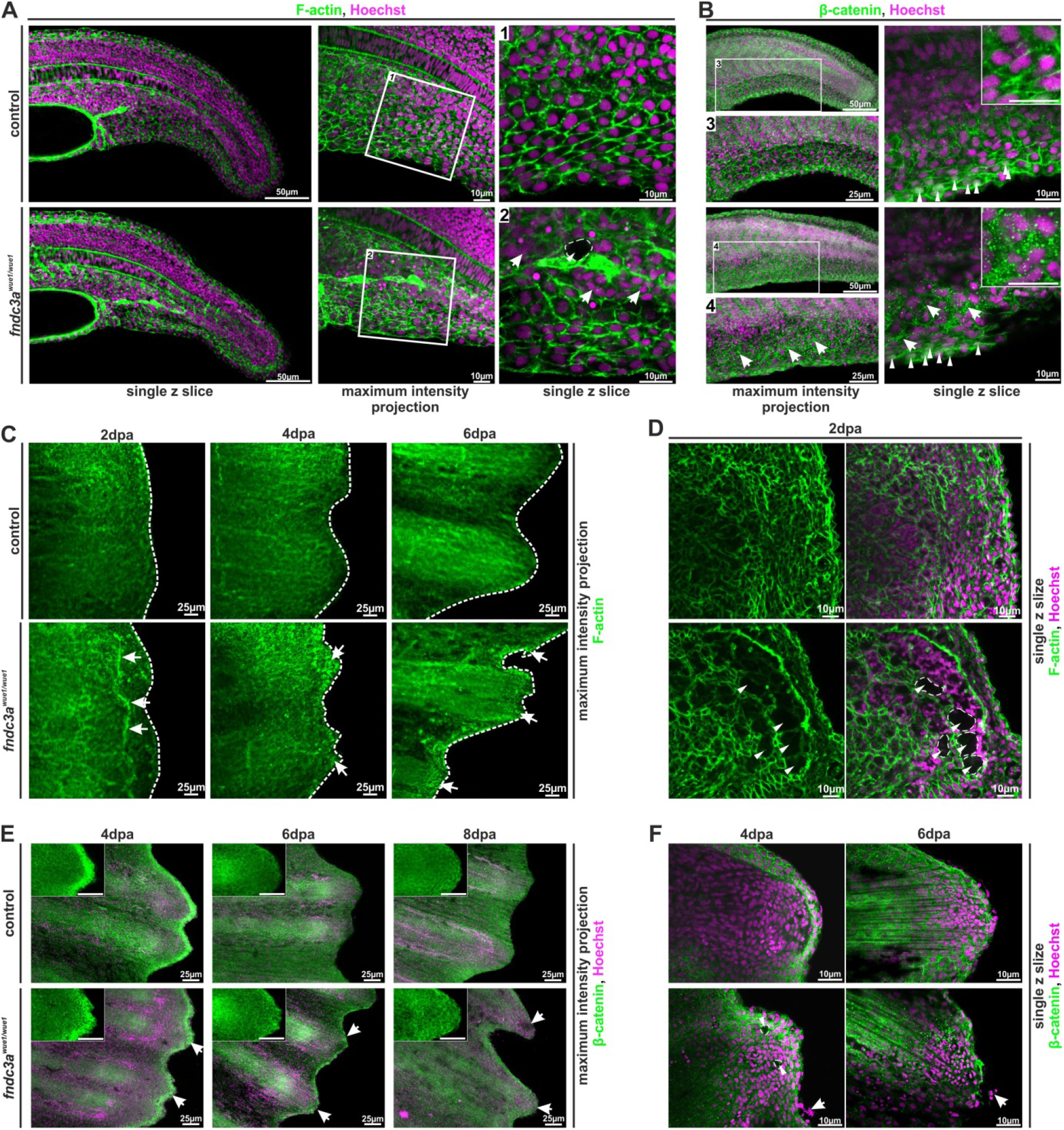
Correct ECM structure in the median fin fold and regenerating caudal fins is hampered in *fndc3a^wue1/wue1^* mutants. (A) F-actin in the median fin fold 22hpf was visualized by phalloidin staining, (B) localization of β-catenin during fin development was visualized by immunofluorescence of 24hpf embryos (n=6 for each group). Both colorations mark cell membranes and ECM structures. Cellular organization of ventral median fin fold cells and ECM matrix is symmetrically structured in control embryos and shows nuclear localization of active Wnt signals in cells at the fin fold tip (white arrowheads in B). *fndc3a^wue1/wue1^* mutants depict cellular alterations and unstructured ECM assembly by showing irregular cell shapes (white arrows in A), cavities within the fin fold (white arrowheads in A2) and speckled accumulation of β-catenin between cells (white arrows in B). Nuclear localization of β-catenin in cells at the fin fold tip was maintained (grey arrowheads in B). (C and D; n=4 for each group) Fin regenerates of *fndc3a^wue1/wue1^* mutants incubated at 32°C and stained for F-actin showed regenerate abnormalities (white arrows in C), irregular regenerate borders (white dashed lines in C) and cellular cavities (white arrowheads in D) (E and F; n=3 for each group) Fin regenerates of *fndc3a^wue1/wue1^* mutants also depicted intracellular accumulation of β-catenin, divergent ECM assembly (white arrows) in addition to appearance of abnormal cells loosely attached to the regenerate (white arrowheads). Images either show maximum intensity projections (30 to 40 single z-slices; z-distance: 1.5µm) or a representative higher resolution single z slice.

## Discussion

Our functional investigations in zebrafish were initiated to proof clinical relevance and the pathomechanism of a novel candidate gene (*FNDC3A*) detected by whole-exome-sequencing in a family diagnosed with split-hand/split-food malformation (SHFM; data not shown). SHFM has been associated with mutations in well described transcription factors, e.g. *DLX5/6* (OMIM *600028; SHFM1; Crackower et al., 1996) or *TP63* (OMIM *603273; SHFM4; Ianakiev et al., 2000). Mouse studies clarified the functional role of TP63 and DLX5 during extremity development in mammals and linked both factors within a common regulatory pathway (Lo Iacono et al., 2008; Restelli et al., 2014). Further investigations show that functional and regulatory analyses of human SHFM candidate genes are feasible in zebrafish and result in detailed insights into molecular function of these genetic factors during vertebrate development (Birnbaum et al., 2012; Heude et al., 2014; Klopocki et al., 2012; Kouwenhoven et al., 2010).

*fndc3a* expression in developing zebrafish was first detected 14hpf in the tail bud and later in apical cells of the ventral median fin fold, the pectoral fins, the notochord and in cells of the chordo neural hinge. Fndc3a protein localization during these early stages of median fin fold development was detected mostly in the cell membrane of ectoderm derived cells and in notochord cells. Spatiotemporally similar expression patterns in the zebrafish median fin fold have been described for other genes causative for SHFM, e.g. *dlx5a* and *tp63* (Bakkers et al., 2002; Heude et al., 2014; Lee and Kimelman, 2002). Comparison of zebrafish *fndc3a* expression during extremity development to mouse *Fndc3a* expression shows a partially similar pattern, i.e. early in the AER of the limb bud and later in interdigital regions of feet (unpublished data; Gene expression database; http://www.informatics.jax.org/expression.shtml). Mouse *Fndc3a* loss-of-function knockouts result in male infertility, while defects in limb development were however not described (Obholz et al., 2006).

The CRISPR/Cas9 generated *fndc3a^wue1/wue1^* mutation in zebrafish, presented in our study, resulted in a premature Stop codon and predicted loss of six fibronectin domains in the corresponding protein. The mutation did not lead to complete nonsense mediated decay and a full loss-of-function phenotype, as residual *fndc3a* mRNA expression could be detected and the observed phenotypes are rather weak and partly transient. Morpholino knockdown and RNA overexpression experiments in wild type and in *fndc3a^wue1/wue1^* mutants, respectively, displayed on the one hand a consistent phenotype of the mutant and further a partial rescue of the induced mutation. On the other hand the experiments resulted also in an increased severity after Morpholino and RNA injection into *fndc3a^wue1/wue1^*, implying that the induced mutation does not abolish the function of the gene completely, but rather leads to a reduced function and results in a hypomorphic mutation. Our results cannot exclude a compensation mechanism for fin development and regeneration in *fndc3a^wue1/wue1^* mutants, which after reduction of Fndc3a function would be able to recover cellular functions and ultimately lead to a rather normally developed caudal fin during larval development and in adult fish. Compensation could either be facilitated by residing functional Fndc3a proteins, as indicated by remaining transcripts in mutants, by other fibronectin homologues, or by activation of known signaling networks during fin development and regeneration (Wehner and Weidinger, 2015). The severe tail phenotype of transient double CRISPR injected individuals suggests a stronger phenotype after complete loss of Fndc3a and points towards interference with prominent signaling pathways, e.g. BMP and TGF-Beta signaling. Potential compensating factors might also be determined by looking at other orthologues of the *fndc3* gene family, e.g. *fndc3ba* and *fndc3bb*, but expression as well as function of these genes have not been investigated yet.

Initial mutant examination revealed a specific phenotype in the developing caudal fin of *fndc3a^wue1/wue1^* mutants, by showing loss of ventral median fin fold cells, loss of gene expression domains in the ventral fin fold, loss of actinotrichia fibers and Col2a accumulation. Most prominently a number of early markers in the ventral median fin fold 20 to 24hpf were reduced or altered in *fndc3a^wue1/wue1^* mutants, indicating cellular Fndc3a function in this specific region and phase of development, which correspond to regions with *fndc3a* expression. In contrast to more globally expressed genes linked to early caudal fin development and to actinotrichia deposition in zebrafish, e.g. *pinfin* mutants lacking Fras1 function or *nagel* mutants lacking Hmcn1 function (Carney et al., 2010), the hypomorphic phenotype in *fndc3a^wue1/wue1^* mutants is milder and locally restricted. One cellular consequence of Fndc3a reduction was suspected to be involved with the assembly process of actinotrichia and collagen fibers during development, as indicated by altered Col2a location in *fndc3a^wue1/wue1^* mutants and by EM imaging. Collagen helix assembly is performed at the ER and is controlled by chaperones, especially Hsp47/SerpinH1 (Ricard-Blum, 2011). Interestingly, knockdown of *hsp47/serpinH1* results in a phenotype resembling *fndc3a^wue1/wue1^*, i.e. actinotrichia organization failure and regenerating fin deformations (Bhadra and Iovine, 2015). In comparison to published loss-of-function mutations in zebrafish collagen genes (e.g. *col1a1*: chi, *Chihuahua; col9a1: prp, persistent plexus*) which result in characteristic severe bone growth defects, skeletal dysplasia and vascular plexus formation (Fisher et al., 2003; Huang et al., 2009) the *fndc3a^wue1/wue1^* mutation has only a mild effect on late caudal fin development by interference with collagen assembly. Additional interference with expression domains of mesenchymal markers, e.g. *myoD*, hints to the notion that Fncd3a is required for the establishment of ventral cells fates of the developing median fin fold, by setting up the correct cellular structure in this region.

The process of median fin fold development is tightly regulated by modulation of epidermal cell shape and correct ECM assembly (Dane and Tucker, 1985; Nagendran et al., 2015). TP63 positive epidermal cells are still present in the *fndc3a^wue1/wue1^* mutants during median fin fold development 20 to 48hpf, but show reduced epidermal cell numbers at ventral positions, delayed ventral fin fold growth and altered morphological properties. Fndc3a shares fibronectin domain III protein domains with other well studied factors like Fibronectin 1, which are known to interact with prominent ECM proteins (Akiyama, 1996) and are essential for matrix assembly (Singh et al., 2010). The observed breakup of correct epidermal cell assembly and appearance of cavities in the ECM of basal epidermal cells after reduction of Fndc3a function indicate that Fndc3a might play a similar role in correct ECM assembly and establishment of correct cellular structures in the median fin fold. Potential downstream effects like misplaced mesodermal cells, loss of actinotrichia fibers and detached epidermal cells are most likely explained by deranged ECM structure in the early epidermal cell layer after Fndc3a reduction. Irregular caudal fin structures, for example, might be a consequence of early developmental irregularities as mesenchymal cells migrate after their induction along predetermined ECM structures and actinotrichia fibers to form the fin skeleton (Feitosa et al., 2012). Future experiments will have to clarify if binding of Fndc3a to integrins is abandoned in *fndc3a^wue1/wue1^* mutants, thereby directly influencing cell adhesion, basal epithelium establishment, and signaling (Towers and Tickle, 2009; Wada, 2011). This idea is supported by the phenotypic similarity of *fndc3a^wue1/wue1^* to *lmna5* and *itga3* mutants (Carney et al., 2010; Nagendran et al., 2015; Webb et al., 2007) and the occurrence of aberrant cells in the epidermal layer of regenerates in the *fndc3a^wue1/wue1^* mutants. Moreover our experiments do not rule out a potential function of Fndc3a during intracellular processes in the Golgi apparatus. Protein localization by (Carrouel et al. 2008) clearly detected FNDC3A in the Golgi apparatus of human odontoblast. Reduction of Fndc3a function therefore may also result in hampered protein processing within this organelle and thereby interfere with modification of ECM proteins or membranous export.

Besides median fin fold development we investigated a potential function of Fndc3a during caudal fin regeneration. In accordance with median fin fold development, regeneration and blastema formation depends on correct epidermal cell assembly and a distinct ECM structure (Govindan and Iovine, 2015; Govindan et al., 2016). We initially detected *fndc3a* transcripts and Fndc3a protein in the distal blastema and in epidermal cell layers of regenerates 2 to 8dpa. In *fndc3a^wue1/wue1^* mutants we observed several temporal effects on caudal fin regeneration: detached epidermal cells, Col2a accumulation, disorganized epidermal cells layers and lacking actinotrichia. These are comparable to the observed developmental phenotype within the median fin fold and indicate ECM malformation to be causative for this effect on adult structures as well.

**Fig.7:**
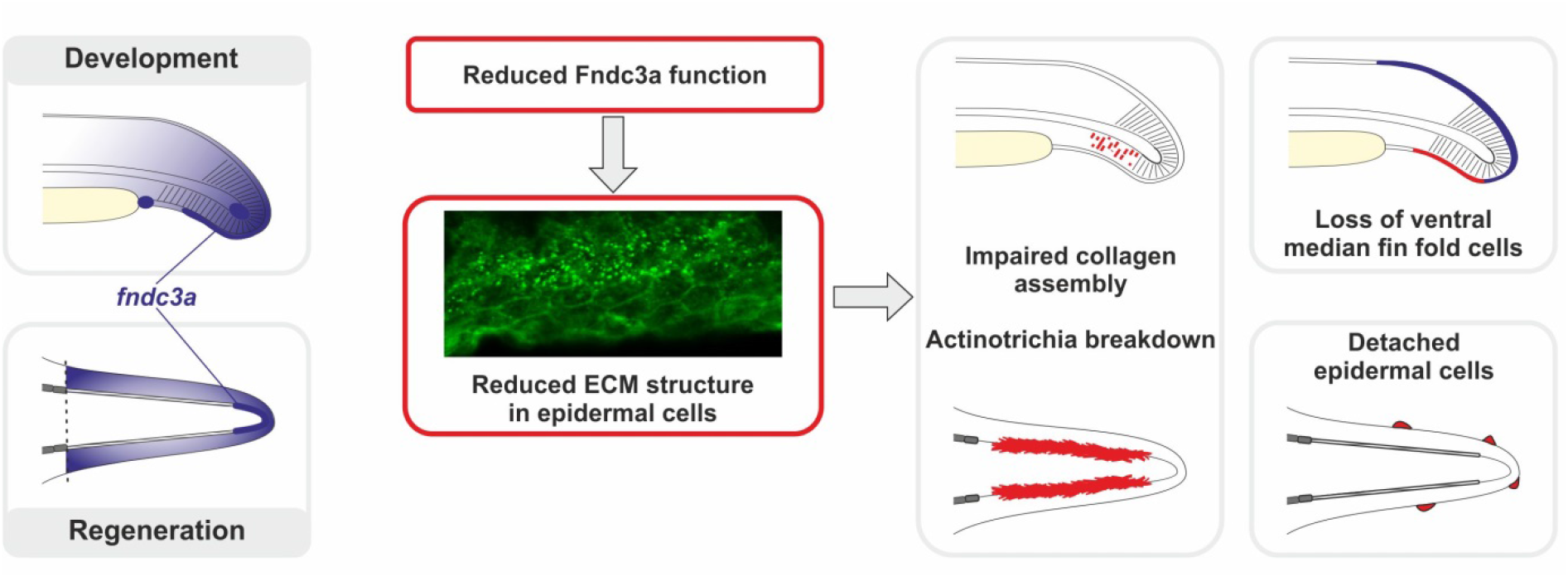
Model of Fndc3a function during zebrafish median fin fold development and caudal fin regeneration. Expression and localization of *fndc3a* can be detected during early phases of median fin fold development and in caudal fin regenerates. Reduced Fndc3a function results in prominent changes of ECM structure of epidermal cells. During development this results in impaired collagen assembly of actinotrichia fibers and loss of ventral median fin fold cells. During regeneration altered ECM structure results in actinotrichia breakdown and detached epidermal cells.

In summary, investigation of *fndc3a* expression and function in zebrafish reveals a transient and spatially restricted role of this genetic factor during extremity development and during fin regeneration. The observed effects after reduction of Fndc3a function on actinotrichia fibers and correct fin morphology are probably secondary and provoked by cellular changes. Most likely disruption of correct ECM structure in basal epidermal cells is the consequential underlying cellular mechanism responsible for the observed fin malformations. Our results demonstrate a cellular link between median fin fold development and caudal fin regeneration due to the necessity for correct cell shape and tissue cohesion in both processes via Fndc3a. Beyond this, our zebrafish experiments now suggest that Fndc3a can influence TP63 positive epidermal cells by altering cell shape or cell adhesion during extremity development. Thus, Fndc3a can be functionally linked to known SHFM genes supporting a potential pathogenic relevance in SHFM phenotypes.

## Materials and Methods

### Animal maintenance

Laboratory zebrafish embryos (*Danio rerio*) of the *AB/TU and AB/AB* strain (ZDB-GENO-010924-10; ZDB-GENO-960809-7) were maintained as described by Westerfield, 2000) under standard aquatic conditions at an average of 24°C water temperature. Embryos were staged by morphological characteristics according to Kimmel et al., 1995). hpf indicate hours-post fertilization at 28.5°C. All procedures involving experimental animals were performed in compliance with German animal welfare laws, guidelines, and policies. Generation of *fndc3a^wue1/wue1^* mutants and fin clipping was approved by the Committee on the Ethics of Animal Experiments of the University of Würzburg and the “Regierung von Unterfranken” (Permit Number: DMS-2532-2-13 and DMS-2532-2-9). The generated *fndc3a^wue1/wue1^* line was submitted to ZFIN.org (ZFIN ID: ZDB-ALT-170417-3).

### CRISPR/Cas9 system and construction of short guiding RNA (sgRNA) constructs

For oligo cloning of sgRNA target sequences the previously published pDR274 vector was used (Hwang et al., 2013) (Addgene Plasmid #42250; sequences of primers used in this study are given in Table S1). For *cas9* RNA synthesis the MLM3613 (Hwang et al., 2013) (Addgene Plasmid #42251) or the pCS2-nCas9n vector (Jao et al., 2013) (Addgene Plasmid #47929) were utilized. Both vectors were purchased from Addgene (www.addgene.org; Cambridge, USA). For designing and constructing of sgRNAs the open access ZiFit Targeter software (http://zifit.partners.org/ZiFiT/) was used. Specific target sites were identified by alignments of zebrafish and human sequences. sgRNA target site
(GGATTCCAGGCCAGTTATGA) is located in exon 13 of *fndc3a* (ENSEMBL Zv9 Transcript: ENSDART00000097261) and targets the second Fibronectin type III domain, while sgRNA target site in exon 18 (GGCGTACAGTGGTTCGGCTC) targets the third Fibronectin type III domain.

### sgRNA transcription and microinjection

sgRNAs were transcribed via the MAXIscript T7 kit (Ambion/ life technologies, Darmstadt, Germany) and were purified via phenol/chloroform extraction. *Cas9* RNA was transcribed via the mMESSAGE mMACHINE kit (Ambion/ Life Technologies, Darmstadt, Germany) and subsequently cleaned via RNeasy purification kit (Qiagen, Venlo, Netherlands). If the MLM3613 vector was used for Cas9 Synthesis, polyA tail synthesis was performed with *E.coli* Poly(A) Polymerase (New England Biolabs, Ipswich, MA, USA) prior to purification. One cell stage zebrafish embryos were injected with solutions comprising sgRNA (25-50ng/µl each), *cas9* RNA (75-100ng/µl), Phenol red (pH7.0; 0.05% final concentration; for visualization of injection solution) and Fluorescein isothiocyanate-dextran (Sigma-Aldrich; 1mg/µl). Positively injected embryos were identified 24hpf by transient green fluorescence of Fluorescein isothiocyanate-dextran and were raised for line establishment.

### Whole mount RNA in-situ hybridization

RNA in situ hybridization was performed according to standard protocols (Hauptmann and Gerster, 1994; Thisse and Thisse, 2008). RNA probes were synthesized from cloned partial mRNA sequences of target genes using the DIG or FLU RNA Labeling Kit (Roche, Basel, Switzerland). We used a *fndc3a* cDNA fragment of 595bp size (used primers: zf_fndc3a_ribo_fwd2 and zf_fndc3a_ribo_rev2) to synthesize a specific anti-sense RNA probe. Sense probes were synthesized as negative control for each anti-sense probe and were used under the same reaction conditions. Primers used for probe cloning are listed in Table S1.

### Immunofluorescence and histology

Immunofluorescence was performed on embryos, cryosections or on whole regenerating fins by standard protocols (Duran et al., 2011; Inoue and Wittbrodt, 2011). Apoptosis assays were performed according to Sorrells et al. (2013) (Sorrells et al., 2013) by visualization of cleaved caspase 3 via immunofluorescence. Primary antibodies used: Fndc3a (HPA008927; Sigma-Aldrich; dilution 1:50-100; Antibody Registry: AB_1078899), Col2a (II-II6B3 was deposited to the DSHB by Linsenmayer, T.F. (DSHB Hybridoma Product II-II6B3); dilution 1:500; Antibody Registry: AB_528165), cleaved Caspase-3 (C92-605; Thermo Fisher Scientific; dilution 1:500; Antibody Registry: AB_397274), β-catenin (610153; BD Biosciences; dilution 1:250; Antibody Registry: AB_397554), TP63 (ab735; Abcam; dilution 1:200; Antibody Registry: AB_305870). Except Fndc3a, all used antibodies have been used previously for experiments in zebrafish and validated protocols have been previously published. Specificity of the used Fndc3a antibody was validated by the Human Protein Atlas project for usage in human tissues (https://www.proteinatlas.org/ENSG00000102531-FNDC3A/antibody). Validation for usage in zebrafish was performed by epitope sequence analysis, antibody dilution series, and incorporation of adequate negative controls in all experiments. Cell nuclei were counterstained with Hoechst 33258 (Sigma-Aldrich; dilution 1:5000). F-Actin was stained with Actin-stain 488 phalloidin (Cytoskeleton, Inc.; PHDG1-A; dilution 1:20).

Whole mount alcian blue staining was performed according to Walker and Kimmel (Walker and Kimmel, 2007). H&E staining on 10μm paraffin sections was performed according to a general histology protocol using Mayer’s Hematoxylin and Eosin.

### Fin regeneration experiments

For fin amputation experiments adult zebrafish (mixed sex, age 3-5 months, 3 independent experiments) were anesthetized with tricaine (MS-222; 3-amino benzoic acid ethyl ester; final concentration ~150mg/l) prior to amputation of ~20% length the of caudal fin. Images were taken 0, 2, 6, 8 and 10 days post-amputation (dpa) under a stereomicroscope. To increase the expected phenotype fish were kept at an increased temperature (32°C) during the experiments. Regenerate samples for immunofluorescence, in-situ hybridization, genotyping, or histology were taken by a second amputation anterior to the first lesion site (2, 4, 6 and 8dpa).

### Image acquisition and quantification

Images were acquired depending on the experiment either with a Leica S8 APO Stereomicroscope (whole embryos), a Zeiss Imager A1 (in-situ hybridizations) or a Nikon A1+ Laser scanning confocal microscope (Immunofluorescence). Image acquisition was performed via device specific cameras/detector and corresponding software (Leica Application suite; Zeiss AxioVision; Nikon NIS-Elements). Further image analyses and quantifications were performed with ImageJ/Fiji (https://fiji.sc/). For figure arrangement CorelDraw Graphics Suite x7 software (Corel Corporation) was used.

Quantification of median fin fold width was performed with ImageJ/Fiji by length measurement of maximum intensity projections of TP63/Hoechst stained embryos 24 and 48hpf. Each embryo was measured at three independent positions at the dorsal or ventral median fin fold. Median fin fold width was measured between the apical fin and somite borders. Quantification of epidermal cells in 24hpf embryos was performed by automated counting of TP63 positive cells in confocal stacks (size z: 40; z-step: 2μm) on the complete ventral and dorsal median fin folds. While in 48hpf embryos three independent positions of the same size (ventral and dorsal median fin fold, tail tip; 8800 μm^2^) were investigated per sample. Used ImageJ plugins and commands: *Stack/Z Project, Despeckle, Watersheed, Analyze particles* (size 2-10μm, circularity 0.5-1.0). A minimum of 12 samples for each condition was investigated.

### Electron microscopy

Embryos or regenerates were washed with PBS and fixed (2.5% glutaraldehyde, 50 mM cacodylat pH 7.2, 50 mM KCl, and 2.5 mM MgCl_2_) overnight at 4°C. Subsequently, embryos or tissues were washed five times with 50 mM cacodyl buffer (pH 7.2) and fixed for 4 h with 2% OsO_4_ in 50 mM cacodylat (pH 7.2) buffer. Embryos or tissues were subsequently stained with 2% uranylacetate overnight. After gradual dehydration with ethanol, they were transferred to propylenoxid and embedded in Epon (SERVA Electrophoresis GmbH, Heidelberg, Germany). Ultrathin sections were analyzed using an EM10 from Zeiss (Oberkochen, Germany).

### qPCR experiments

For qPCR experiments RNA was extracted from pools of 12 embryos each (age 24hpf; F3 *fndc3a^wue1/^+* and *fndc3a^wue1/wue1^* generation; control: AB). For each genotype three independent pools were generated and compared. For cDNA synthesis 1μg RNA was transcribed into cDNA and further analyzed in a ViiA7 Real-Time PCR System (Thermo Fisher Scientific). Each group and primer sample was analyzed in triplicates on a single qPCR plate utilizing HOT FIREPol Eva Green Mix Plus (Solis Biodyne). Data analysis of was performed via QuantStudio Real-Time PCS Software v1.1 by ΔΔCt method. *AB* control samples were used as reference sample for relative comparison. Amplification of *gapdh* and *eef1a1l1* was used as endogenous/housekeeping controls. Used primer pairs, amplicon sizes and targeted regions are noted in table S1. Experiments were performed according to MIQE guidelines.

## Acknowledgements

The authors thank Anja Kirschbauer for professional fish keeping support, Tabea Röder for in-situ probe synthesis, Prof. Dr. Manfred Schartl for sharing aquatic equipment, as well as Claudia Gehrig and Daniela Bunsen from the Electron Microscopy Unit of the Biozentrum at the University of Würzburg, for preparation of ultrathin sections and support with the electron microscopy analysis. We are indebted to Prof. Dr. Christoph Winkler for fruitful discussions.

## Data availability

## Author contributions

D.L., S.G. and E.K. designed the experiments. D.L., I.K., S.K., M.M. and M.O. conducted the experiments. D.L. and E.K. wrote the manuscript with input from all authors.

